# Commensal acidification of specific gut regions produces a protective priority effect against enteropathogenic bacterial infection

**DOI:** 10.1101/2025.02.12.637843

**Authors:** Jane L. Yang, Haolong Zhu, Puru Sadh, Kevin Aumiller, Zehra T. Guvener, William B. Ludington

## Abstract

The commensal microbiome has been shown to protect against newly introduced enteric pathogens in multiple host species, a phenomenon known as a priority effect. Multiple mechanisms can contribute to this protective priority effect, including antimicrobial compounds, nutrient competition, and pH changes. In *Drosophila melanogaster*, *Lactiplantibacillus plantarum* has been shown to protect against enteric pathogens. However, the strains of *L. plantarum* studied were derived from laboratory flies or non-fly environments and have been found to be unstable colonizers of the fly gut that mainly reside on the food. To study the priority effect using a naturally occurring microbial relationship, we isolated a wild-fly derived strain of *L. plantarum* that stably colonizes the fly gut in conjunction with a common enteric pathogen, *Serratia marcescens*. Flies stably associated with the *L. plantarum* strain were more resilient to oral *Serratia marcescens* infection as seen by longer lifespan and lower *S. marcescens* load in the gut. Through *in vitro* experiments, we found that *L. plantarum* inhibits *S. marcescens* growth due to acidification. We used gut imaging with pH-indicator dyes to show that *L. plantarum* reduces the gut pH to levels that restrict *S. marcescens* growth *in vivo*. In flies colonized with *L. plantarum* prior to *S. marcescens* infection, *L. plantarum* and *S. marcescens* are spatially segregated in the gut and *S. marcescens* is less abundant where *L. plantarum* heavily colonizes, indicating that acidification of specific gut regions is a mechanism of a protective priority effect.

## Introduction

The animal gut is colonized by a stable microbiome that is specific to its host species(1–4), but transient bacteria also colonize and contribute a large part of the microbiome diversity(5, 6). There are host mechanisms for acquiring and maintaining the stable bacteria as well as for filtering the transient colonizers and excluding pathogens(7–9). These host mechanisms involve the immune system and metabolic processes such as the production of stomach acids and bile salts(7). Stable microbes in the gut can also prevent invasion by pathogens through what are known as ecological priority effects(10–12). Priority effects play a crucial role during ecological succession and are defined as processes in which established colonizers influence the colonization of newly arriving species(11, 13).

In the gut microbiome, studies have shown that symbiotic bacteria competitively exclude pathogens from establishing themselves in numerous host species(14–19). Several potential mechanisms are thought to drive this process, including competition for space(20, 21), competition for nutrients(22), production of inhibitory compounds such as antibiotics(23), and changing the physiology of the gut environment(24–26) including by acidifying the gut lumen through production of short chain fatty acids (27–29); these components are difficult to differentiate in a complex community because there are many indirect effects, which are challenging to experimentally separate. The use of gnotobiotic animals greatly reduces the experimental complexity.

*Drosophila* has emerged as a model microbiome system due to its naturally low complexity of approximately 5 core species of intestinal bacteria, all of which are culturable(30). Lactobacilli form a major component of the species diversity in flies as well as in humans and include multiple species that are known to be beneficial probiotics for their hosts(31–33).

Lactobacilli form biofilms(20), crossfeed with other bacteria(34), and are known to produce reuterin and other antimicrobials that inhibit pathogens(35–37). Lactobacilli also metabolize carbohydrates to produce lactic acid(34, 38–41), which acidifies their environment to a pH of ∼4 and can inhibit the growth of other species(41, 42). *Lactiplantibacillus plantarum* is widely prevalent across multiple animal guts and is one of the most widely studied human probiotics(43–45). It is also one of the most common species of bacteria in Drosophila guts and has been shown to have probiotic properties such as enhancing fly nutrition(33, 46, 47), stimulating immunity(48), and preventing colonization by enteric pathogens(19, 29).

*Serratia marcescens* is a common fly and human enteric pathogen found in both wild and laboratory environments(15, 49, 50). Studies in flies have found that *L. plantarum* reduces the pathogenesis of *Serratia*(19), yet the mechanism remains unclear. Moreover, the majority of *Drosophila* microbiome studies use strains of bacteria that have been isolated from laboratory flies(30), which often show signatures of evolution to the laboratory environment and may have lost some of their natural traits(51–53).

Here we isolate *L. plantarum* and *S. marcescens* from a single wild fly and study their effects on fly health and interactions with each other. We find that *L. plantarum*’s acidification of the gut plays a significant role in the inhibition of *S. marcescens* and the resulting benefit to the fly.

## Results

### *Pre-colonization with* L. plantarum *increases fly survival in response to* S. marcescens *infection*

Different strains of the same bacterial species may have a wide range of properties; wild isolates can behave differently from lab strains in terms of virulence(54–56) and colonization ability(57). In order to study bacteria that stably colonize the fly gut, we caught a wild *D. melanogaster* female, passaged it daily to fresh, germ-free food for 12 days to clear out transient colonizers, then surface sterilized it with 70% ethanol before crushing and plating to isolate colonies. The fly was confirmed to be *D. melanogaster* by Sanger sequencing the *cytochrome oxidase I* gene. We isolated 76 colonies with 5 distinct morphologies (**Table 1**). Each strain was passaged to single colonies 5 consecutive times on streak plates and freezer stocks were made. We then Sanger sequenced the 16S genes of 20 of these isolates, representing all 5 morphologies, to confirm that each was a single species isolate. We found 7 commensal species, *L. plantarum*, *Levilactibacillus brevis*, *Acetobacter orientalis*, *A. tropicalis*, *A. cerevisiae*, *A. malorum*, and *A. sicerae*, as well as *S. marcescens*, an opportunistic pathogen (**Table 1**).

**Table 1.**
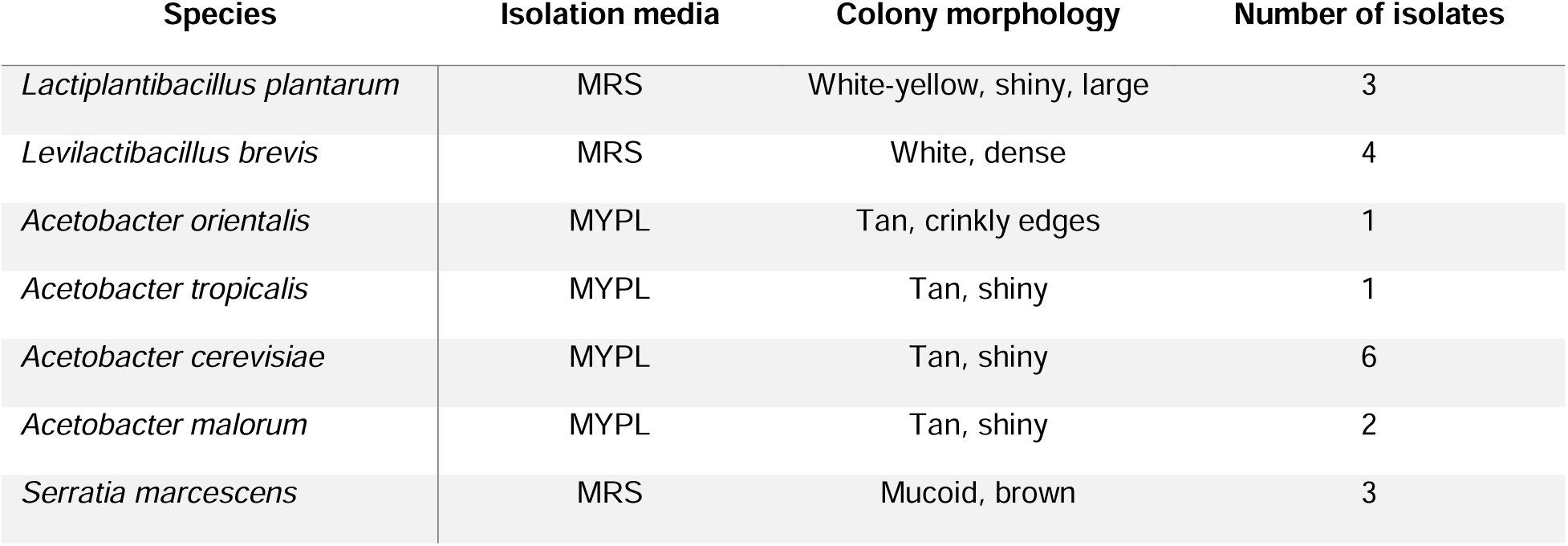
Bacterial strains isolated from an individual wild *D. melanogaster*.

It has been previously shown that *L. plantarum* isolated from lab flies has a protective effect against *S. marcescens* infection (19). Using our wild-fly strains of the two bacteria, we tested whether *L. plantarum* is protective against *S. marcescens* oral infection. We started with germ-free, 5-day-post eclosion mated female flies. One group of flies was kept germ-free; a second group was colonized with *L. plantarum* for 3 days; a third group was orally infected with *S. marcescens* after 3 days of pre-colonization by *L. plantarum*; a fourth group was orally infected with *S. marcescens* without pre-colonization with *L. plantarum*. We then measured the lifespan of each treatment group of flies (Fig. 1A). Germ-free flies lived the longest (median 45 days), followed by *L. plantarum*-colonized (median 37 days). Flies colonized by *S. marcescens* lived only 17 days. Pre-colonization with *L. plantarum* increased the median lifespan to 27 days, a 59% increase (**Fig. 1A,B**). Co-colonizing flies simultaneously with *L. plantarum* and *S. marcescens* (median lifespan of 15 days) did not produce the increase in lifespan (**Fig. 1C,D**), indicating that *L. plantarum* must be established first. These results confirm that *L. plantarum* from wild flies has a protective effect on flies orally infected with *S. marcescens*.

**Figure 1.**
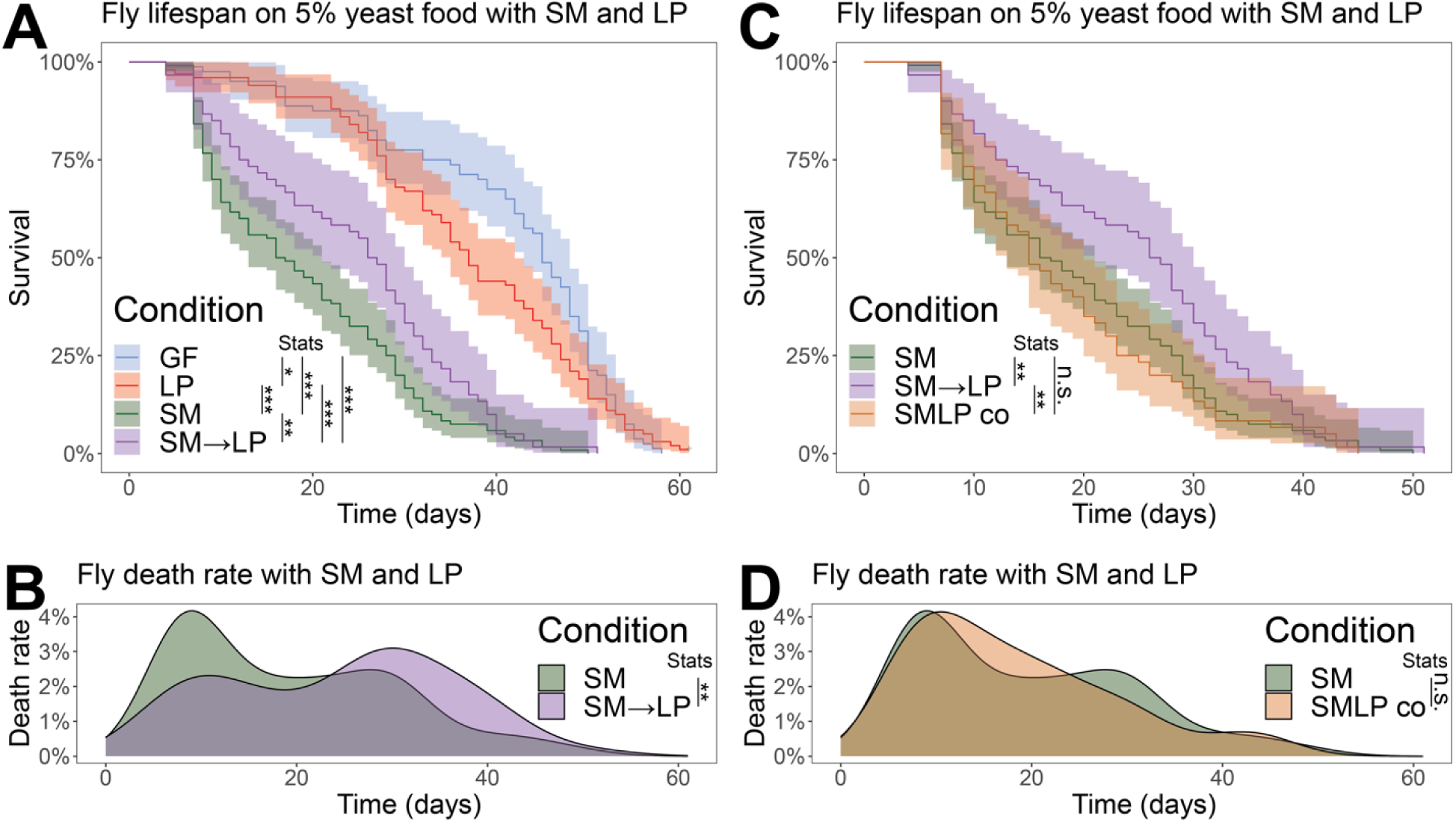
Pre-colonization with *L. plantarum* increases fly survival in response to *S. marcescens* infection. (A) Lifespans of flies with different microbial inoculations, germ-free (GF, blue), colonized by *L. plantarum* (LP, red), colonized by *S. marcescens* (SM, green), and colonized by *L. plantarum* then *S. marcescens* sequentially (SM→LP, purple). (B) Data for SM and SM→LP plotted as death rates. (C) Lifespan data comparing SM and SM→LP verus flies co-inoculated with SM and LP at the same time (SMLP co, orange). (D) Data in C plotted as death rates. N=80 GF; N=100 LP; N=120 SM; N=60 SM→LP; N=60 SMLP co. 3 or more separate biological replicates per treatment. Kruskil-Wallis ANOVA with Wilcoxon rank sum post hoc with Benjamini & Hochberg correction.

### L. plantarum *colonization diminishes* S. marcescens *abundance in the* Drosophila *gut*

We hypothesized that the protective effects of L. plantarum are due to a decrease in S. marcescens abundance in the presence of L. plantarum. To test, we inoculated flies as before for the lifespan experiments and then sacrificed flies at 4-day intervals over the next 16 days to enumerate colony forming units (CFUs) of each bacterial strain. In flies inoculated with a single species of bacteria, the L. plantarum abundance remained constant at ∼10^5^ CFUs per fly over the 16 days (**Fig. 2A**), while the S. marcescens abundance increased over the first 8 days before stabilizing at ∼10^5^ CFUs per fly (**Fig. 2B**). However, in flies pre-colonized with L. plantarum before S. marcescens inoculation, the S. marcescens abundance was an order of magnitude lower at ∼10^4^ CFUs per fly (**Fig. 2B**), confirming our hypothesis that L. plantarum decreases S. marcescens colonization of the fly gut in a protective priority effect. To determine whether the effect occurs on the fly food versus within the gut, we plated L. plantarum and S. marcescens on agar plates with the fly food composition. No growth of S. marcescens and minimal growth of L. plantarum was observed on fly food after 4 days at 30°C, indicating that the inhibitory effects we observed occur primarily inside the fly.

**Figure 2.**
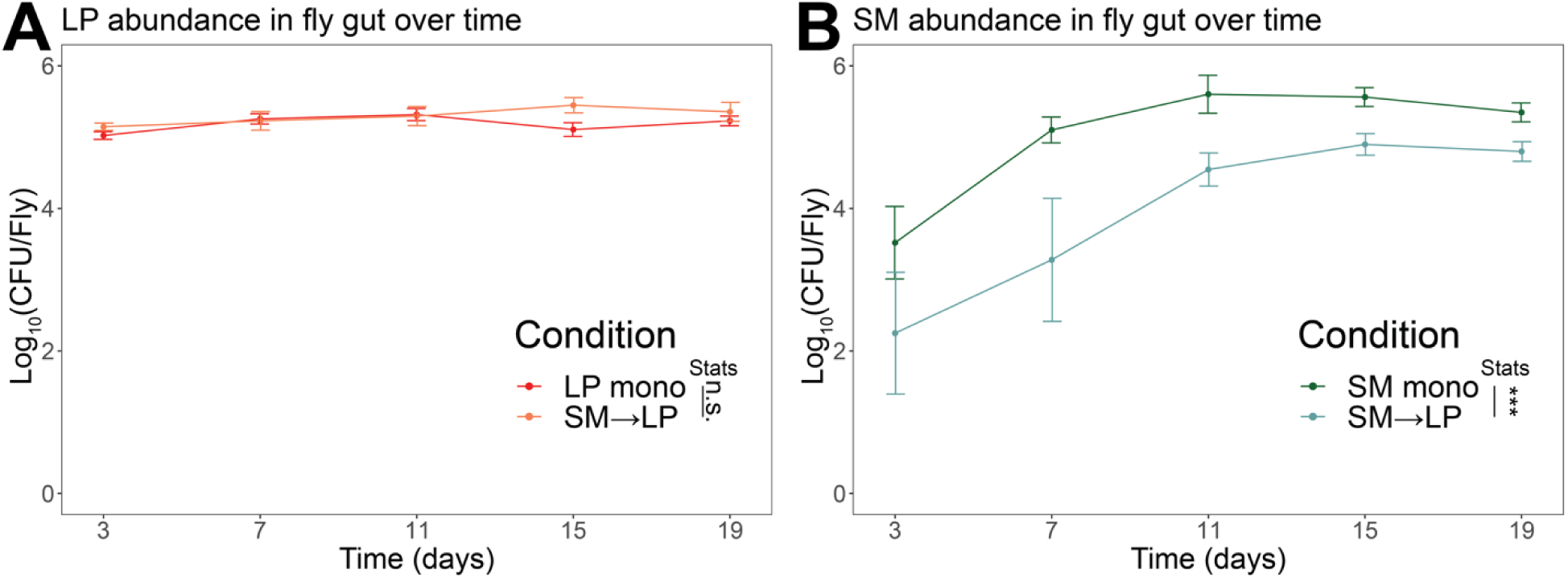
*L. plantarum* colonization diminishes the abundance of *S. marcescens* in the *Drosophila* gut. (A) *L. plantarum* abundance in the fly gut in colony-forming units (CFUs) comparing *L. plantarum* mono-colonized flies (LP mono, N=40) with flies sequentially colonized by *L. plantarum* then *S. marcescens* (SM→LP, N=37). (B) *S. marcescens* abundance in the fly gut comparing *S. marcescens* mono-colonized flies (SM mono, N=40) with flies sequentially colonized by *L. plantarum* then *S. marcescens* (SM→LP, N=37). Wilcoxon rank sum test.

### L. plantarum *reduces* S. marcescens *growth through acidification*

We next tested whether *L. plantarum* inhibits the growth of *S. marcescens* in culture in liquid MRS media. Consistent with the abundances inside the fly, co-cultures in liquid media decreased *S. marcesens* growth by ∼1 order of magnitude (**Fig. 3A-B**). To investigate the inhibitory mechanism, we evaluated four hypotheses; (i) *L. plantarum* inhibits *S. marcescens* through contact-dependent effects, (ii) *L. plantarum* produces an antimicrobial compound, (iii) *L. plantarum* utilizes nutrients required by *S. marcescens*, and (iv) *L. plantarum* acidifies the growth environment.

**Figure 3.**
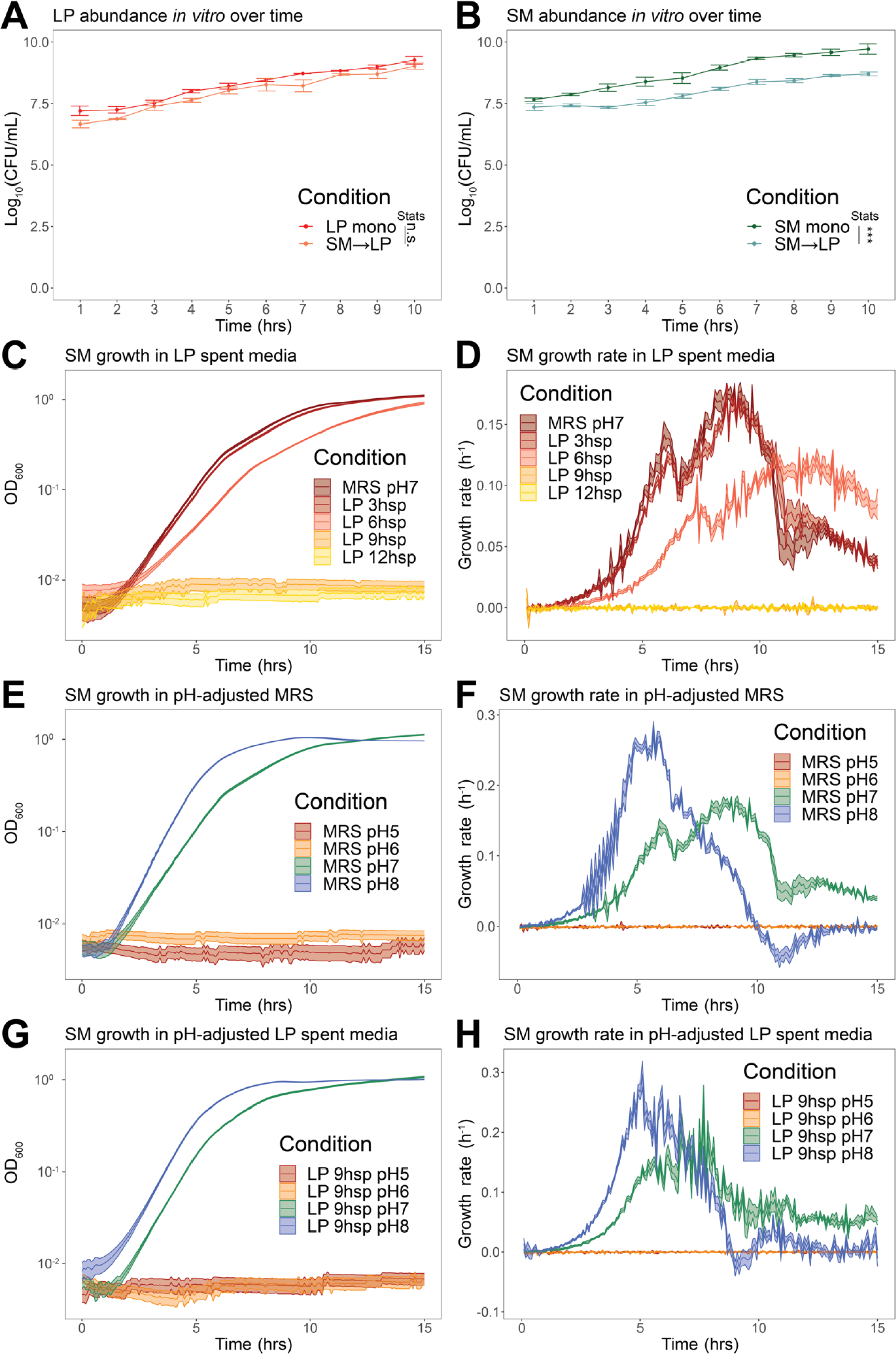
*L. plantarum* reduces *S. marcescens* growth through acidification. (A) *In vitro L. plantarum* abundance with (N=29) or without (N=30) *S. marcescens* in co-culture as assessed by CFU counts. (B) *In vitro S. marcescens* abundance with (N=29) or without (N=30) *L. plantarum* in co-culture as assessed by CFU counts. (C) Growth of *S. marcescens* in spent media of *L. plantarum* as assessed by optical density at 600 nm (OD_600_). (D) Growth rates of the cultures in C plotted over time. (E) Growth of *S. marcescens* in MRS media of different pH in the range of 5 to 8. (F) Growth rates of cultures in E plotted over time. (G) Growth of *S. marcescens* in *L. plantarum* 9 hour spent medium with pH adjusted in the range of 5 to 8. (H) Growth rates of cultures in G plotted over time. N=18 per growth curve; 3 biological replicates and 6 experimental replicates per biological replicate. Wilcoxon rank sum test for (A) and (B).

We first tested whether spent media from *L. plantarum* impacts the growth of *S. marcescens*. We grew *L. plantarum* for 3 hours, 6 hours, 9 hours, and 12 hours in liquid MRS media then removed *L. plantarum* cells by centrifugation followed by filtration through a 0.22 µm PES membrane. We then inoculated *S. marcescens* into these spent media preparations and assayed growth by optical density at 600 nm (OD_600_) in a microplate spectrophotometer. *S. marcescens* showed slight growth impairment in 3-hour spent media, significant impairment in 6-hour spent media, and no growth in 9-hour or 12-hour spent media (**Fig. 3C**), indicating that contact-dependent effects are not necessary for the observed inhibition of *S. marcescens*. More specifically, the growth rate of *S. marcescens* is lower in spent media in which *L. plantarum* has grown for a longer period of time. (**Fig. 3D**).

We determined that the pH of the *L. plantarum* spent media was below pH 5 by 12 hours of culture, which is consistent with previous work reporting detailed pH measurements over time(40). For this reason, we suspected that acidification of the media by *L. plantarum* inhibited *S. marcescens* growth. We next measured the pH-dependence of *S. marcescens* growth in MRS. *S. marcescens* showed optimal growth at pH 8, impaired growth at pH 7, and no growth at pH 6 or below (**Fig. 3E**), indicating that the pH of the *L. plantarum* spent media is sufficient to cause the observed growth inhibition of *S. marcescens*. As seen with the spent media, these differences were due to changes in growth rate (**Fig. 3F**) rather than the length of lag phase.

While we found that pH changes are sufficient to cause the observed growth inhibition, we could not rule out that *L. plantarum* produces a compound that directly inhibits *S. marcescens* growth. To test this hypothesis, we increased the pH of 9-hour *L. plantarum* spent MRS medium to see if this would restore *S. marcescens* growth. We found that adjusting the pH to ≥7 restored *S. marcescens* growth (**Fig. 3G**) and growth rate (**Fig. 3H**) to similar levels as fresh MRS. This suggests that there is no inhibitory compound present in the spent media, and pH changes alone are sufficient to cause the observed growth inhibition. In addition, the nutrition depletion by *L. plantarum* over 9 hours had no significant effect on *S. marcescens* growth.

In evaluating our four initial hypotheses, a contact-dependent effect is not supported because the *S. marcescens* inhibition occurs in the absence of any physical contact with *L. plantarum*. An antimicrobial compound is not a primary means of the inhibition because *S. marcescens* growth is normal in spent medium with raised pH. Nutrient competition is also an unlikely mechanism, because *S. marcescens* is able to grow in media that has been nutritionally depleted by *L. plantarum*. Thus, *L. plantarum* acidification of the growth medium is the main process inhibiting *S. marcescens* growth *in vitro*.

### L. plantarum *acidifies the Drosophila foregut, hindgut and rectum*

Based on our *in vitro* experiments, we predicted that *L. plantarum* growth acidifies the fly gut, helping to limit *S. marcescens* growth. To measure the regional pH throughout the gut, we fed flies pH indicator dyes. Bromocresol green can indicate pH differences in the range of ∼3 to 5. Phenol red can indicate pH differences in the range of ∼6 to 8. We prepared flies as before, either keeping them germ-free or inoculating with *L. plantarum*, *S. marcescens*, or both bacteria. We then allowed the flies to adjust for 3 days, dissected their guts, and imaged. To quantify the pH, we prepared standards and imaged them simultaneously with the gut samples.

The copper cell region of the midgut showed a pH of ∼4.5 in all flies regardless of microbial treatment (**Fig. 4**), consistent with other studies(58). Similarly, the anterior midgut had a pH of ∼7 in all flies. Differences in pH corresponding to the different microbial treatments were observed in the foregut, posterior midgut, hindgut, and rectum. In germ-free flies, the crop had a pH of ∼7, whereas it was ∼4 in *S. marcescens*-treated flies and ∼3.5 in *L. plantarum*-treated and *L. plantarum+S.marcescens*-treated flies. The posterior midgut had a pH of >8 in germ-free flies, 7.9 in *S. marcescens*-treated flies, 7.6 in *L. plantarum*-treated flies, and 7.2 in *L. plantarum+S.marcescens*-treated flies, suggesting minor acidification by *L. plantarum* in this region.

**Figure 4.**
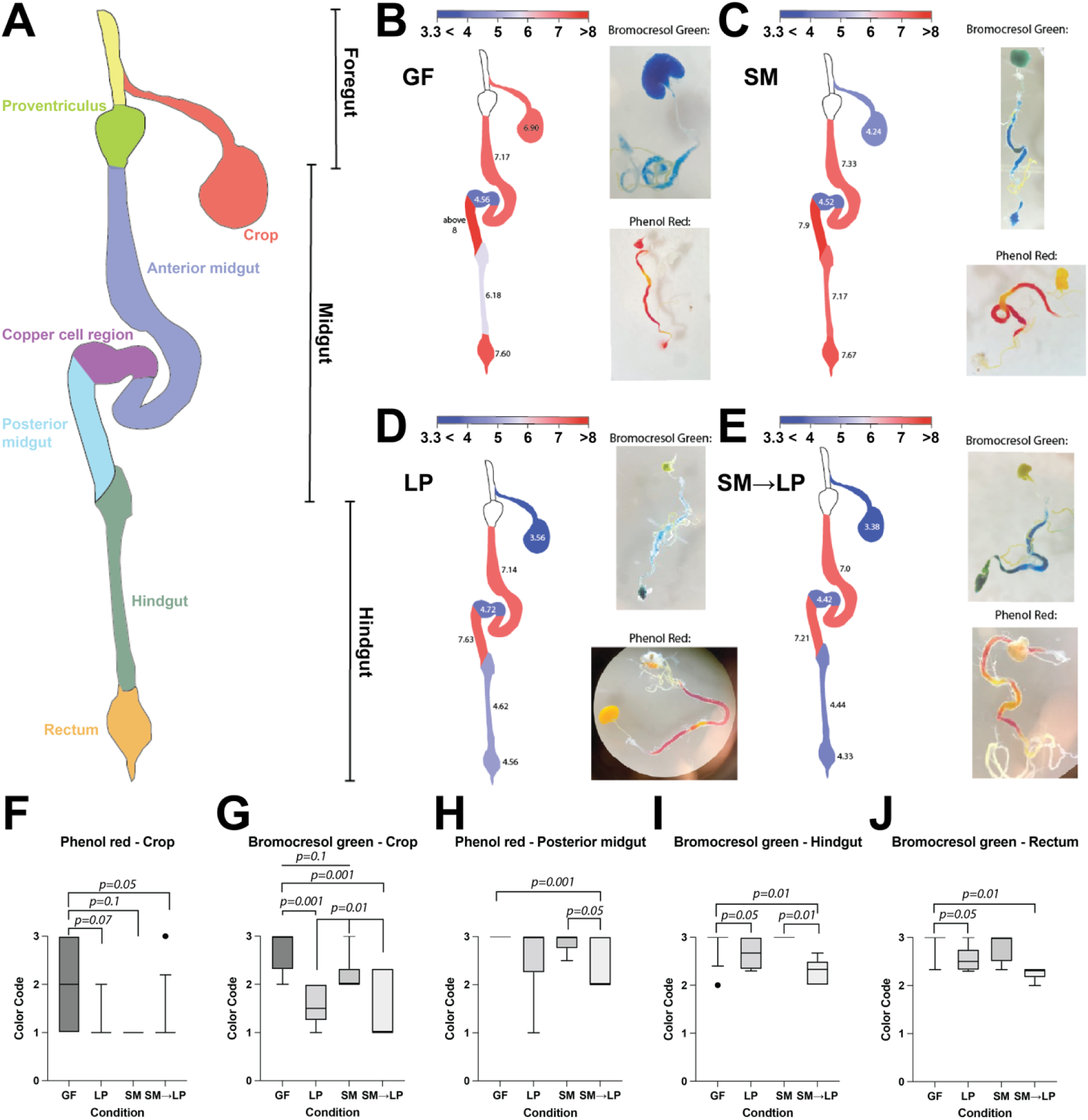
*L. plantarum* acidifies the foregut, hindgut and rectum. (A) Diagram of *Drosophila* gut. (B-E) pH indicator dyes bromocresol green and phenol red show the pH of each region of the gut in flies that are (B) germ-free (GF), (C) colonized by *S. marcescens* (SM), (D) colonized by *L. plantarum* (LP), or (E) colonized by *L. plantarum* then *S. marcescens* (SM→LP). Numbers on gut regions indicate average pH measured. (F-J) statistical comparison of pH differences between the microbial treatments for the crop, posterior midgut, hindgut, and rectum using a color code. Note that the entire midgut region’s pH is consistent regardless of bacterial colonization. N=76 total flies quantified. For phenol red, N=10 GF, N=8 LP, N=6 SM, N=13 SM→LP. For bromophenol green, N=13 GF, N=9 LP, N=7 SM, N=10 SM→LP. Wilcoxon rank sum test with Holm-Bonferroni correction.

The largest differences were observed in the hindgut and rectum. In germ-free flies, the hindgut pH was ∼6. In *S. marcescens*-colonized flies, the hindgut pH was ∼7, while in both *L. plantarum*-treated flies and *L. plantarum+S.marcescens*-treated flies, the hindgut pH was ∼4.5. Similarly, the rectum pH was ∼7.5 in germ-free and *S. marcescens*-colonized flies, while it was ∼4.5 in *L. plantarum*-treated and *L. plantarum+S.marcescens*-treated flies. Across all gut regions sampled, flies with both *L. plantarum* and *S. marcescens* were more similar in pH to flies inoculated with *L. plantarum* than the other treatment groups. Overall, our results indicate that *L. plantarum* colonization acidifies the foregut, hindgut and rectum but has only minor effects on the midgut pH.

### S. marcescens *does not co-localize with* L. plantarum *in the gut*

To test whether the low pH gut regions have reduced *S. marcescens* abundance, we used plasmids to make fluorescently labeled strains of *L. plantarum* (mCherry) and *S. marcescens* (mGFP), colonized flies with these strains, dissected the guts, and imaged to determine the spatial localization (**Fig. 5**). *L. plantarum* was present in all regions of the gut, including the crop, anterior midgut, posterior midgut, hindgut, and rectum with varied abundances in the different regions. *S. marcescens* was predominantly found in the anterior midgut, consistent with the hypothesis that it cannot tolerate low pH regions of the gut.

**Figure 5.**
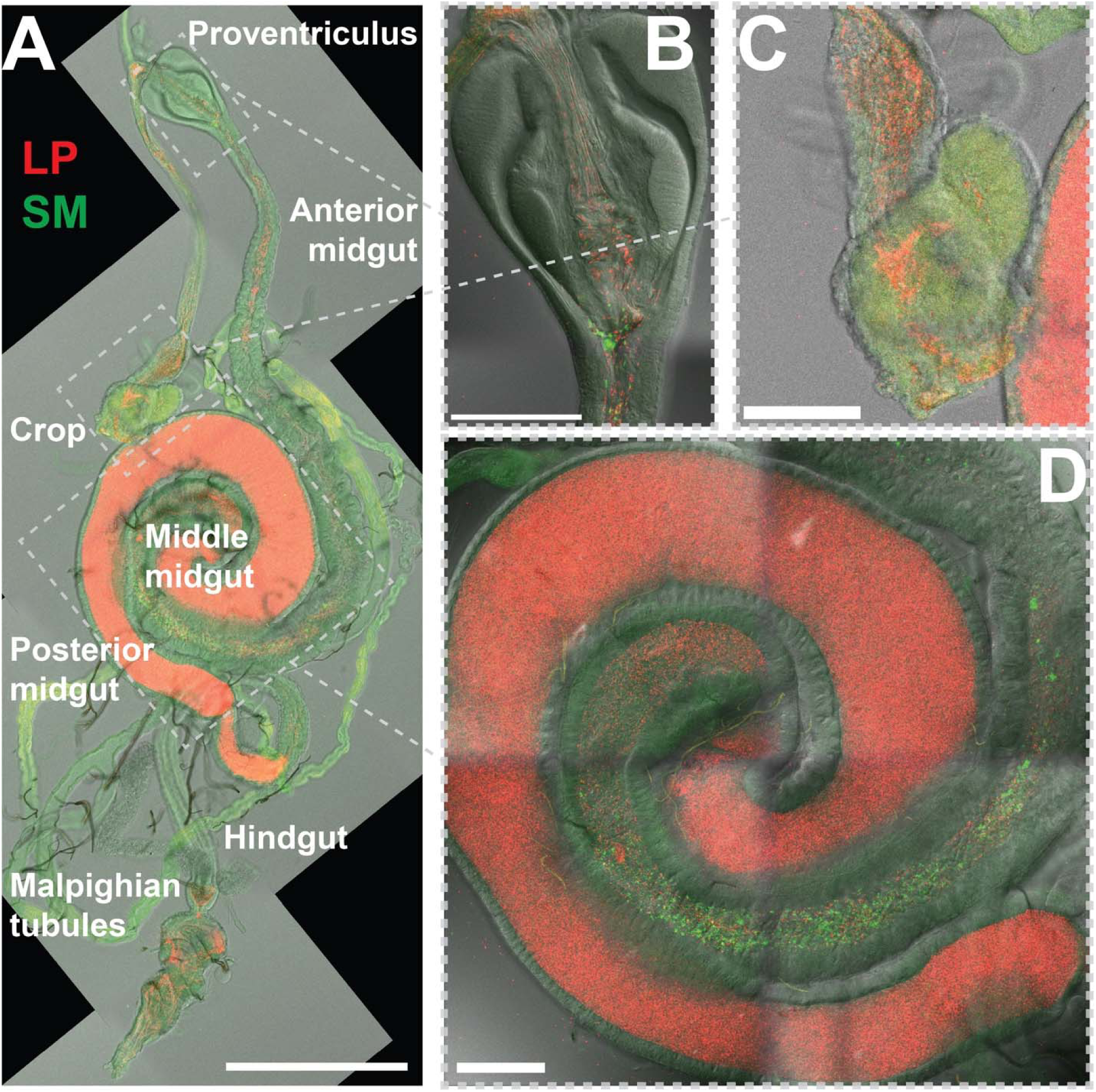
*S. marcescens* does not co-localize with *L. plantarum* in the gut. (A) Fluorescence micrograph of *Drosophila* gut co-colonized by *L. plantarum* (mCherry-labeled, red) and *S. marcescens* (mGFP-labeled, green, scale bar is 500 µm). (B-D) Closeups of (B) proventriculus, (C) crop, and (D) middle midgut with copper cell region, showing spatial segregation of *L. plantarum* and *S. marcescens* (scale bars are 100 µm). Note that green autofluorescence from the fly tissue in A prevents accurate visual localization of *S. marcescens*. Green puncta in B and D are *S. marcescens*. Representative images from a single gut. N=8 guts from 3 biological replicates.

## Discussion

### Synopsis

We found that *L. plantarum* prevents pathogenesis of *S. marcescens* through a priority effect in the gut. While indirect mechanisms, such as stimulation of host immunity or mucus production, can mediate protective effects of the gut microbiome, direct bacteria-bacteria inhibition can also occur through a variety of mechanisms, including contact-dependent inhibition, resource competition, production of antibacterial secondary metabolites, and changing the pH of the environment. Through *in vitro* and *in vivo* studies, we found that acidification drives *L. plantarum* inhibition of *S. marcescens* growth inside the fly.

### Wild fly versus lab fly strains

Because wild fly gut communities differ from those in the lab(59, 60), we isolated bacteria from healthy wild flies to examine the process by which lactobacilli species protect the host from infection. We found that wild-fly associated *L. plantarum* thrives in the gut and improves host survival during chronic *S. marcescens* infection (**Fig. 1**). At the same time, the wild fly strain of *S. marcescens* was able to survive in the acidic gut environment and maintain a stable, albeit small population in the fly gut (**Fig. 2**) despite an inability to grow on the fly food. Unlike some strains of *S. marcescens*, the strain we isolated did not cause immediate death and instead formed a chronic infection that shortened the fly’s lifespan.

We note that this strain of *L. plantarum* is distinct from another strain that we isolated from a different wild-caught fly. The latter strain we have found to form large, long-term biofilms in the foregut(57), but the strain studied here does not form the same dense aggregates in the foregut. Future studies could examine the effects of *L. plantarum* biofilm formation on *S. marcescens* establishment to infer the role of spatial structure on pathogen exclusion.

### Priority effects and coexistence

We were able to distinguish the spatial location of the priority effects as occurring in the host gut rather than on the food. The lack of growth on the food is likely because the food pH is too acidic (pH 4.5) for either bacterial species to grow(40, 41), and the sugar content (10% glucose) is inhibitory due to the high osmolarity(61, 62). By imaging the fly gut with different pH indicators, we found that the gut pH is lower when *L. plantarum* is present, particularly in the crop, hindgut and rectum regions (**Fig. 4**). We also found that while *L. plantarum* inhabits the entire gut, *S. marcescens* is mostly limited to anterior midgut regions where the pH is higher and *L. plantarum* density is lower. The spatial segregation of these two wild-fly derived species may allow them to coexist within the gut. The low *S. marcescens* population may limit virulence within the host, allowing flies to survive for a longer period.

*L. plantarum* is most likely the source of the acidification that inhibits *S. marcescens*. *D. melanogaster* produces an acidic region in its midgut, called the copper cell region, the cells of which are conserved with mammalian copper cells in the stomach, which produce the digestive stomach acids(58, 63). Outside of the copper cell region, the fly gut pH of germ-free flies is close to neutral. However, multiple regions of the gut acidify when *L. plantarum* colonizes. No known fly cell types exist in the crop, posterior midgut, hindgut, or rectum that produce acid, whereas *L. plantarum* cells are definitive acid producers. Thus, the only likely mechanism of the acid production in the crop, hindgut, and rectum is through *L. plantarum*, which clearly inhabits those gut regions and produces acid.

We note that the highest abundance of *L. plantarum* appeared in the posterior midgut, a region that was not acidic in any of the conditions. The midgut regulates important digestive processes including the breakdown and absorption of food and therefore requires tighter pH regulation to control these enzymatic and cellular processes. We suggest that the midgut, which is endoderm-derived tissue, may regulate pH in the posterior midgut in addition to the copper cell region. By contrast, the foregut, hindgut and rectum are ectoderm-derived tissue and covered by intestinal cuticle, which may make pH more difficult for the host to regulate in those regions, yet the resident bacteria in the foregut, hindgut and rectum may be more competent in regulating pH by secreting acidic molecules such as lactic acid.

We also note that the crop pH decreased to ∼4 in *S. marcescens* mono-colonized flies, which we speculate is due to low pH food in the crop because *S. marcescens* does not significantly lower the pH. Colonization by *L. plantarum* or *L. plantarum*+*S. marcescens* lowered the pH to ∼3.5, which is lower than the food and indicates the metabolic activity of *L. plantarum*.

### Acidification as a mechanism of microbe-microbe interactions in the gut

Our study shows that *L. plantarum* can diminish the negative effects of chronic enteric pathogen infection by inhibiting the pathogen through acidification of certain gut regions, which complements recent work showing that *Drosophila* gut microbiome acidity on the food protects flies from the fly pathogens *Pseudomonas entomophila* and *Erwinia carotovora* by production of lactic acid(29). Prior studies have shown that acidification is a strong driver of microbe-microbe interactions *in vitro* (41, 42, 64). A specific investigation of the interactions between *L. plantarum* and *S. marcescens* showed bistability of co-cultures, where either *L. plantarum* or *S. marcescens* would drive the other species extinct during long-term passaging(41). Consistent with those results, we found that initial colonization with *L. plantarum* was necessary for the inhibition of *S. marcescens* in the fly gut and the subsequent benefits for fly lifespan (**Fig. 1**). In contrast, we found long-term stable coexistence of *L. plantarum* and *S. marcescens* in the fly gut (**Fig. 2**). We suggest that this coexistence may be supported by the spatial segregation of the two species (**Fig. 5**), consistent with environmental heterogeneity of the gut tissues promoting diversity of the microbiome.

## Acknowledgements

1. G. Lopez (Synvivia) generously transformed the *S. marcescens* strain with a GFP-expression plasmid. R. Grabherr and S. Heinl generously provided the pCD256-mCherry expression plasmid for *L. plantarum*. W.B.L. acknowledges NIH grants DP5OD017851, R01DK128454, and R21AI173779 as well as NSF grants IOS 2032985 and IOS 2144342.

## Materials and methods

### Drosophila husbandry

*Drosophila melanogaster* Canton-S strain was the standard laboratory wild-type for the study. Flies used were germ-free adult mated females. Fly food was made of 10% glucose, 5% autoclaved live yeast, 1.2% autoclaved agar, and 0.42% propionic acid, all sterilized. The food has a pH of 4.5. Fly stocks were transferred to fresh food every 3-4 days.

### Bacteria inoculation and fly lifespan measurement

*L. plantarum* (LP) and *S. marcescens* (SM), both isolated from a single wild *D. melanogaster*, were used in the study. Bacteria cultures were prepared in liquid MRS at 30°C overnight. Bacteria were orally introduced to the flies by spreading ∼10^5^ CFU of bacteria suspended in 50 μL of MRS across the surface of sterile fly food which subsequently housed 20 germ-free flies per vial. Flies were inoculated with LP on day 3-4 and SM on day 6-7 by allowing the flies to feed on inoculation media for 24 hours. Control groups were fed only one of the two bacteria strains or kept germ-free. All flies were then transferred onto sterile media and flipped into new, sterile vials every 3-4 days for the remainder of the experiment. Fly survival was recorded daily. A Kruskil-Wallis ANOVA with a Wilcoxon rank sum post hoc test was performed in R with a Benjamini & Hochberg correction.

### Fly gut bacterial load quantification

Flies were reared and inoculated as described earlier. Every 3-4 days following SM inoculation, 7-8 female flies were randomly selected from each treatment for bacterial load quantification. Flies were anesthetized under CO_2_, and then washed in 70% ethanol and PBS for surface sterilization. Next, a mechanical pestle was used to homogenize each fly individually in 1 mL PBS. The suspensions were then diluted, plated onto MRS agar plates, and left to grow for 40 hours at 30°C. Colony morphology was used to identify and count CFUs. A Wilcoxon rank sum test was performed in R.

### Bacteria *in vitro* coculture

SM and LP were grown separately in liquid MRS at 30°C. Each strain of bacteria was then introduced into 50 mL of liquid MRS to create two suspensions with a final OD_600_ of 0.01. Coculture was prepared by mixing 25 mL of each suspension. All flasks were left to grow on a continuous shaker at 30°C, and 1 mL of suspension from each flask was taken and plated every hour to monitor the growth of the two bacteria. Plated MRS agar plates were left for incubation at 30°C for 40 hours. CFUs were identified based on morphology and counted. A Wilcoxon rank sum test was performed in R.

### Preparation of LP spent media and pH-adjusted media

LP from a frozen stock was cultured overnight in liquid MRS at 30°C as initial LP inocula. MRS media inoculated with LP at a starting OD_600_ of 0.01 was prepared and left to grow at 30°C. Liquid culture was collected at different time intervals, then centrifuged to pellet cells and the supernatant was vacuum filtered through a 0.22 µm PES membrane. For fresh MRS media and LP spent media requiring pH adjustment, the media pH was adjusted accordingly using diluted HCl and NaOH and measured using pH paper and a pH probe.

### SM *in vitro* growth measurement

SM *in vitro* growth under different media conditions was measured using a BioTek Epoch 2 Microplate Spectrophotometer. Each well of a 96-well plate was prepared with 198 μL of respective media and 2 μL of SM suspension at an OD_600_ of 1 to reach a final OD_600_ of 0.01. SM was cultured at 30°C with shaking and SM growth was monitored by measuring OD_600_ every 5 minutes.

### Fly gut pH measurement

Flies were reared as described earlier on fly food with an addition of 0.1% the respective pH indicator dye. Bromocresol Green (BG) and Phenol Red (PR) were used separately to examine the pH range of 3.33-5.00 and 6.33-8.00 respectively. Bacteria inoculations were prepared as described earlier and 30 μL of each bacterial suspension was spread on the surface of each fly food. A group of 20 flies was transferred to each vial and left to feed for 5 days. Fly gut tissues were dissected under a ZEISS Stemi 508 dissecting microscope. Each piece of fly gut tissue was observed in 200 μL of PBS (pH 7.4). Samples were discarded if color was detected in PBS indicating the leakage of luminal fluid. Fly gut tissues were gently straightened and mounted on slides for image acquisition.

First, to generate pH dye color standards, pH gradients with 0.1% of each of the pH dyes were created and measured in a 96-well plate with pH adjusted PBS using diluted HCl and NaOH and measured using pH paper and a pH probe. Images were captured using the same lighting, imaging setup, and equipment as for the fly gut tissues.

Images were examined using ImageJ. Each image was analyzed by measuring the average RGB values of each of the 6 fly gut regions and then comparing them to the pH dye standards. A color code system was created according to (65) to quantify the pH. For PR, yellow (indicating pH ≤ 6.33) is defined as 1, orange (indicating pH = 7) is defined as 2, and red (indicating pH ≥ 8) is defined as 3. For BG, yellow (indicating pH ≤ 3.33) is defined as 1, green (indicating pH = 4) is defined as 2, and blue (indicating pH ≥ 5) is defined as 3. A Wilcoxon rank sum test was performed in MATLAB with the Holm-Bonferroni correction for multiple comparisons.

### Generation of fluorescently labeled bacteria strains

*L. plantarum* strain ZTG301 was transformed with pCD256-p11-mCherry as described in (66–68). Briefly, overnight cultures were diluted to OD_600_ = 0.1 in MRS media supplemented with 1% glycine and 0.75 M Sorbitol and grown to OD_600_ = 0.4-0.6 under constant shaking. Upon reaching desired densities, cultures were harvested by centrifugation at 4500 x g for 5 minutes at 4°C and washed twice with ½ culture volume ice-cold buffer (0.95 M sucrose, 3.5 mM MgCl2). Final cell suspensions were prepared in 1/25 culture volume buffer supplemented with 10% glycerol and split into 50 µL aliquots in 1.5 mL microcentrifuge tubes. Cell suspensions were flash frozen in liquid nitrogen and stored at -80°C until use. Frozen aliquots were thawed on ice for 10 minutes followed by incubation with 1 µg unmethylated plasmid DNA obtained from C2925 *E. coli* on ice for 10 minutes. Suspensions were then transferred to ice-cold 2 mm electroporation cuvettes and electroporated at 2000 Vm 25 µF, 400 Ohms in a Bio-Rad Genepulser Xcell electroporation system. Transformations were recovered with 950 µL MRS supplemented with 0.5 M sucrose and 0.1 mM MgCl_2_ for 2 h at 37°C, followed by plating onto MRS agar plates supplemented with 10 µg/mL chloramphenicol.

*S. marcescens* strain ZTG300, was transformed with plasmid pCC2828, containing an ampicillin resistance marker and constitutive sfGFP expression construct. For electroporation, we followed protocols for *E. coli*. Briefly, we made electrocompetent cells by growing to and OD_600_ of 0.3. Cells were washed 3 times with ice cold 10% glycerol, and electroporated in 1 mm cuvettes at 1600V, 200 Ohms, and 25 µF. Transformations were recovered with 950 µL SOC medium for 1 h at 37°C, followed by plating onto LB agar plates supplemented with 100 µg/mL ampicillin.

### Bacteria *in vivo* imaging in fly guts

Flies were pre-inoculated with LP for 3 days as described earlier. On the day of dissection, flies were briefly transferred to sterile 1.2% agar vials for 2 hours to stimulate feeding during the subsequent SM inoculation. Flies were then inoculated with SM for 1 hour before being collected for dissection. Fly gut tissues were isolated, processed, and mounted on slides for imaging as previously described(69). Images were captured using a Leica TCS SP8 confocal microscope.

